# Multi-species ELISA for the detection of antibodies against SARS-CoV-2 in animals

**DOI:** 10.1101/2020.08.26.266825

**Authors:** Kerstin Wernike, Andrea Aebischer, Anna Michelitsch, Donata Hoffmann, Conrad Freuling, Anne Balkema-Buschmann, Annika Graaf, Thomas Müller, Nikolaus Osterrieder, Melanie Rissmann, Dennis Rubbenstroth, Jacob Schön, Claudia Schulz, Jakob Trimpert, Lorenz Ulrich, Asisa Volz, Thomas Mettenleiter, Martin Beer

## Abstract

Severe acute respiratory syndrome coronavirus 2 (SARS-CoV-2) has caused a pandemic with millions of infected humans and hundreds of thousands of fatalities. As the novel disease - referred to as COVID-19 - unfolded, occasional anthropozoonotic infections of animals by owners or caretakers were reported in dogs, felid species and farmed mink. Further species were shown to be susceptible under experimental conditions. The extent of natural infections of animals, however, is still largely unknown. Serological methods will be useful tools for tracing SARS-CoV-2 infections in animals once test systems are validated for use in different species. Here, we developed an indirect multi-species ELISA based on the receptor-binding domain (RBD) of SARS-CoV-2. The newly established ELISA was validated using 59 sera of infected or vaccinated animals including ferrets, raccoon dogs, hamsters, rabbits, chickens, cattle and a cat, and a total of 220 antibody-negative sera of the same animal species. Overall, a diagnostic specificity of 100.0% and sensitivity of 98.31% was achieved, and the functionality with every species included in this study could be demonstrated. Hence, a versatile and reliable ELISA protocol was established that enables high-throughput antibody detection in a broad range of animal species, which may be used for outbreak investigations, to assess the seroprevalence in susceptible species or to screen for reservoir or intermediate hosts.

## Introduction

Since the beginning of 2020, the acute respiratory disease COVID-19 caused by a novel betacoronavirus, severe acute respiratory syndrome coronavirus 2 (SARS-CoV-2) (Zhu et al., 2020), has been keeping the world in suspense. COVID-19 emerged for the first time in December 2019 in Wuhan, China (WHO, 2020b), and has developed rapidly into a global pandemic (WHO, 2020a) resulting in millions of infections and hundreds of thousands of deaths. The symptoms in affected humans range from inapparent infections to fever, fatigue, cough, shortness of breath or severe pneumonia and death (WHO, 2020c). SARS-CoV-2 is primarily transmitted between infected humans through droplets and fomites (Ghayda et al., 2020; WHO, 2020c). However, the virus is thought to originate from an animal reservoir, from where it was likely transmitted to humans by either a direct spill-over event presumably followed by natural selection and adaptation in humans, or via an intermediate mammalian host (Andersen, Rambaut, Lipkin, Holmes, & Garry, 2020). As SARS-CoV-2 is highly similar to bat betacoronaviruses (Latinne et al., 2020; Wu et al., 2020; Zhou et al., 2020) or betacoronaviruses found in pangolins (Zhang, Wu, & Zhang, 2020), it is suspected that one of those animal species may represent the original host of the SARS-CoV-2 ancestral virus. Considering the presumed zoonotic origin of SARS-CoV-2 and the fact that highly similar orthologues of the human angiotensin-converting enzyme (ACE2) receptor are present in certain animal species (Li, 2013; Sun, Gu, Ma, & Duan, 2020), it is of interest to identify susceptible animal species and to assess SARS-CoV-2 prevalence in these species. As the COVID-19 pandemic unfolded, occasional infections of dogs, domestic and non-domestic felid species (cat, tiger, lion) and farmed minks were reported (Enserink, 2020; McAloose et al., 2020; Oreshkova et al., 2020; Sailleau et al., 2020; Sit et al., 2020; Wang et al., 2020). The majority of SARS-CoV-2-infections in animals were directly linked to SARS-CoV-2-infected owners or animal caretakers. Under experimental conditions, the susceptibility of additional species including ferrets, hamsters, and fruit bats has been demonstrated (Osterrieder et al., 2020; Richard et al., 2020; Schlottau et al., 2020; Shi et al., 2020). However, the level of natural infections is largely unknown. Currently, infection in animals is mostly confirmed by direct virus detection using (real-time) RT-PCRs. While this is highly sensitive and specific, experimental and field data demonstrate that animals test positive only during a very short time interval. Therefore, to elucidate the host range of SARS-CoV-2 and the prevalence in susceptible species serological assays, i.e. multi-species antibody detection systems, could be beneficial. Serum neutralization tests, which are species-independent, rely on *in-vitro* interaction between the infectious virus and specific antibodies and, therefore, require high-containment laboratories to handle the virus. In contrast, enzyme-linked immunosorbent assays (ELISAs) can be applied under less stringent biosafety conditions, once such test systems are validated for use in animals. In addition, ELISAs enable high-throughput testing of clinical specimens.

For human sera, a wide range of ELISA systems have been developed within a very short time based on different antigens that include the viral nucleocapsid (N) protein, the spike protein (S) or the S receptor-binding domain (RBD) (Beavis et al., 2020; Klumpp-Thomas et al., 2020; Tré-Hardy et al., 2020). When comparing different antigens as assay targets, considerable cross-reactivity between different coronaviruses occurred in whole-virus or N-based serological assays, while S or RBD based protocols demonstrated a much higher specificity (Chia et al., 2020; Klumpp-Thomas et al., 2020; Meyer, Drosten, & Müller, 2014). Based on these findings, we selected different domains of the S protein for the development of a multi-species SARS-CoV-2 antibody ELISA to avoid cross-reactivity and associated non-specificity. This is especially important as coronaviruses including several betacoronaviruses are widespread in animals (Amer, 2018; Drechsler, Alcaraz, Bossong, Collisson, & Diniz, 2011; Erles & Brownlie, 2008; Hodnik, Ježek, & Starič, 2020; Murray, Kiupel, & Maes, 2010; Tizard, 2020), and could potentially cross-react in SARS-CoV-2 test systems.

## Materials and methods

### Protein expression and ELISA procedure

For expression of the SARS-CoV-2 S1 and the RBD-SD1 domain, amino acids (aa) 17 to 685 or 319 to 519, respectively, were amplified from a codon-optimized synthetic gene (GeneArt synthesis, ThermoFisher Scientific, Germany). The constructs were cloned into the pEXPR103 expression vector (iba lifesciences, Germany) in-frame with an N-terminal modified mouse IgΚ light chain signal peptide and a C-terminal double Strep-tag. Expi293 cells were grown in suspension in Expi293 expression medium (ThermoFisher Scientific) at 37 °C, 8 % CO_2_, and 125 rpm. The cells were transfected using the ExpiFectamine293 transfection kit (ThermoFisher Scientific) according to the manufacturer’s instructions. Five to six days after transfection, the supernatants were harvested by centrifugation at 6000xg for 20 min at 4 °C. Biotin was blocked by adding BioLock (iba lifesciences) as recommended, and the supernatant was purified using Strep-Tactin XT Superflow high capacity resin (iba lifesciences) according to the manufacturer’s instructions. Finally, the proteins were eluted with 50 mM biotin (in 100 mM Tris-HCl, 150 mM NaCl, 1 mM EDTA; pH 8.0). The expression of the proteins was verified by SDS-PAGE and western blotting using a HRP-conjugated anti-Strep-tag antibody (dilution 1/20,000, iba lifesciences). Reactive bands were visualized using the Super Signal West Pico Chemiluminescent substrate (ThermoFisher Scientific) and an Intas ChemoCam system (Intas Science Imaging Instruments GmbH, Germany).

For the ELISA, medium-binding plates (Greiner Bio-One GmbH, Germany) were coated with 100 ng/well of either the RBD or S1 antigen overnight at 4 °C in 0.1 M carbonate buffer (1.59 g Na_2_CO_3_ and 2.93 g NaHCO_3_, ad. 1 l aqua dest., pH 9.6), or were treated with the coating buffer only. Thereafter, the plates were blocked for 1 h at 37 °C using 5 % skim milk in phosphate-buffered saline (PBS). To evaluate the optimal concentration, sera were pre-diluted 1/50, 1/100 or 1/200 in Tris-buffered saline with Tween 20 (TBST), and incubated on the coated and uncoated wells for 1 h at room temperature. A multi-species conjugate (SBVMILK; IDvet, France) was diluted 1/40 or 1/80 and then added for 1 h at room temperature. For chicken samples, an anti-chicken conjugate (Rabbit anti-Chicken IgY (H+L) Secondary Antibody, HRP; ThermoFisher Scientific) diluted 1/10,000 was applied. Following the addition of tetramethylbenzidine (TMB) substrate (IDEXX, Switzerland), the ELISA readings were taken at a wavelength of 450 nm on a Tecan Spectra Mini instrument (Tecan Group Ltd, Switzerland). Between each step, the plates were washed three times with TBST. The adsorbance was calculated by subtracting the optical density (OD) measured on the uncoated wells from the values obtained from the protein-coated wells for the respective sample.

### Serum samples

A total of 220 animal sera (51 cat, 33 cattle, 32 ferret, 27 raccoon dog, 35 chicken, 30 rabbit, 12 hamster) collected before the SARS-CoV-2 pandemic or representing pre-infection sera of SARS-CoV-2 animal studies were included as negative control samples. The SARS-CoV-2 negative status of the latter was verified by a virus neutralization test (VNT) or in an indirect immunofluorescence assay (iIFA).

Eight ferret, 20 hamster, 16 raccoon dog and two cattle SARS-CoV-2 antibody-positive sera originated from experimental infection studies and were collected between 8 and 28 days post infection (dpi) (Freuling et al., 2020; Osterrieder et al., 2020; Schlottau et al., 2020; Ulrich, Wernike, Hoffmann, Mettenleiter, & Beer, 2020; Balkema-Buschmann, Rissmann et al., unpublished). With the exception of the bovine samples, all sera tested positive in a VNT, in an iIFA, or in both, with the iIFA being more sensitive for antibody detection early after infection (Schlottau et al., 2020). The cattle sera were collected 12 and 20 dpi from the same animal. The first sample tested negative by iIFA, while the second showed weak positive signals (titer 1/4). In addition, three sequential sera from a naturally SARS-CoV-2 infected cat were included (ProMED-mail, 2020). The titers of the feline samples as measured by VNT were 1/40 (serum collected 15 days after the first SARS-CoV-2 positive throat swab sample), 1/51 (22 days) and 1/40 (29 days), respectively. Furthermore, sera of three chickens (49 days post prime and 28 days post boost immunization, iIFA positive >1/640) and two rabbits immunized with the RBD and three chickens (49 days post prime and 28 days post boost immunization, iIFA inconclusive) and two rabbits immunized for immuno-serum generation with the S1 protein were added.

### Sensitivity, specificity, repeatability and reproducibility

To determine the cut-off values and the diagnostic sensitivity and specificity of the final ELISA protocol, the sera were tested, and receiver operating characteristic (ROC) analyses were performed.

For evaluation of intra-assay reproducibility, a negative and a very weak-positive cattle serum (collected at 12 dpi) as well as a positive raccoon dog serum (16 dpi) were tested in five replicates each. The inter-assay repeatability was determined with the identical samples and replicate numbers on five independent ELISA plates. Mean values and standard deviations of the 25 replicates were calculated. All analyses and visualizations were performed using GraphPad Prism version 8.0 for Windows (GraphPad Software, USA).

## Results and Discussion

Both SARS-CoV-2 protein domains, i.e. RBD and S1, were successfully expressed (figure 1), and their functionality was proven by their reactivity with sera from infected or immunized animals (figures 2 and 3). Thereafter, both proteins were compared regarding the optimal conditions for their use as capture antigens in antibody ELISAs. In general, a higher absorption signal was obtained for positive control sera when the RBD domain was coated, independent of the applied serum or conjugate concentration, while S1 as target antigen led to a stronger background signal for negative control samples in every approach (figure 2). It has been reported in previous studies that the RBD domain is the most specific antigen for the detection of SARS-CoV-2 antibodies in humans, predominantly because this region is a major target of antibodies, and, in addition, is poorly conserved between different coronaviruses (Amanat et al., 2020; Chia et al., 2020; Premkumar et al., 2020). Considering the results of the RBD/S1 comparison in this study, the same holds true for animals. Alternatively, the superior performance of the RBD domain might be related to the protein production and purification procedure, a phenomenon that has been already described for SARS-CoV-2 proteins (Klumpp-Thomas et al., 2020). While the expression of the RBD is relatively straightforward, the production of the large-size, heavily glycosylated S1 domain is more challenging. Both, expression and purification are inefficient and consequently require pooling of several independently produced batches and more hands-on-time, which might lead to a lower quality of the final protein preparation. Here, we selected the RBD domain for further use because of the stronger specific signal combined with a lower unspecific background signal.

**Figure 1:**
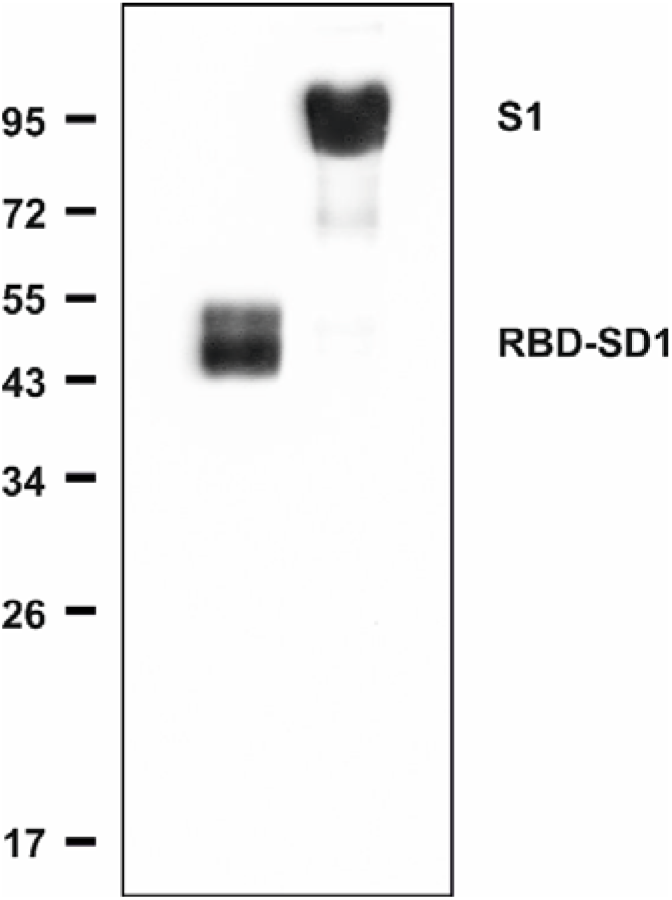
Western blot analyses of purified RBD-SD1 and S1 proteins used for plate coating and/or immunization.

**Figure 2:**
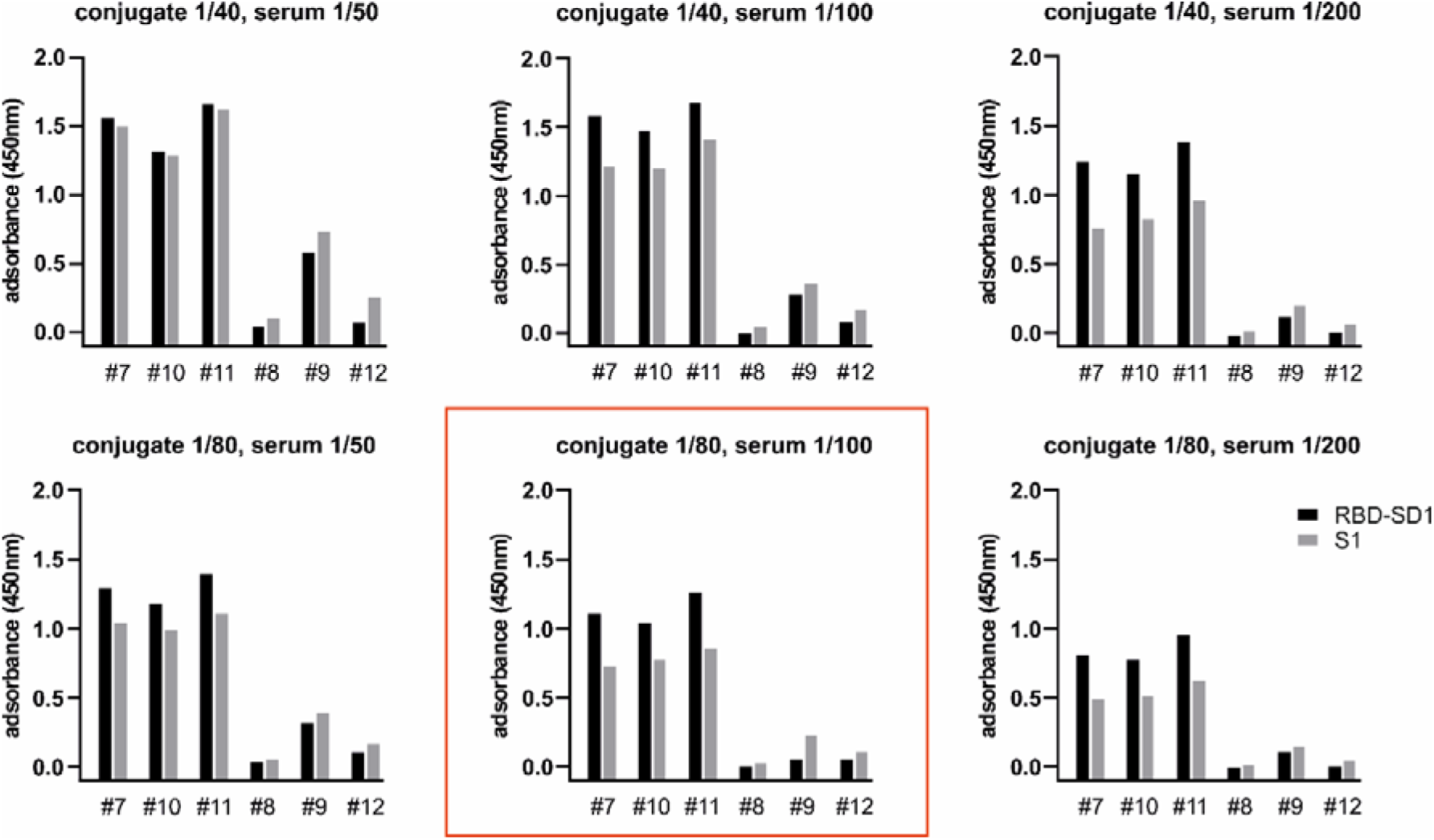
Optimization of the SARS-CoV-2 ELISA. Three raccoon dog sera that tested positive by an indirect immunofluorescence assay (animal numbers #7, #10 and #11) and three negative sera (#8, #9 and #12) were diluted 1/50, 1/100 or 1/200, and tested in combination with a multi-species conjugate (dilution 1/40 or 1/80) against the SARS-CoV-2 RBD-SD1 domain (black bars) or S1 protein (grey bars) expressed in Expi293 cells. The serum-conjugate combination that was selected for the final ELISA protocol is framed in red.

**Figure 3:**
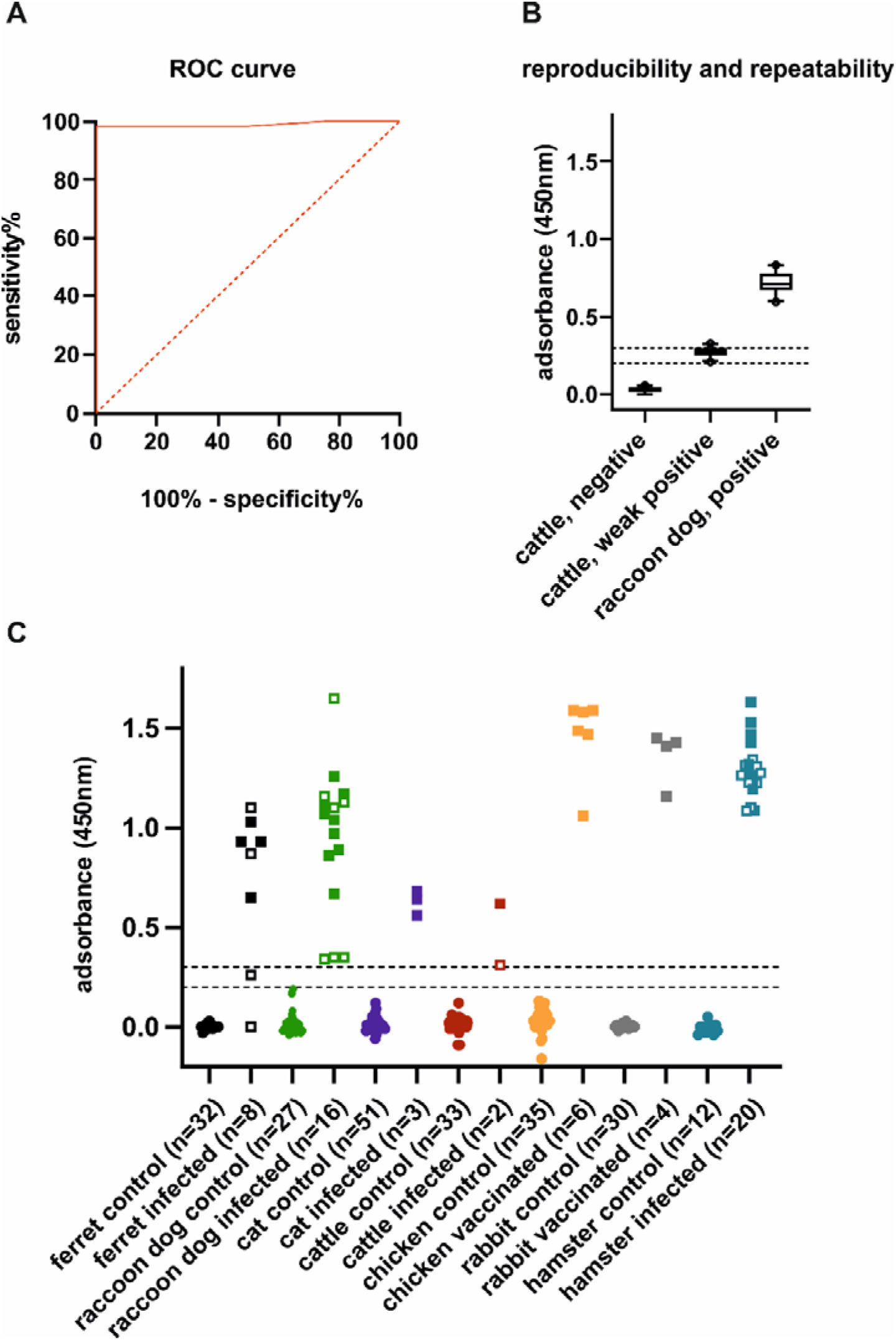
Assay validation and diagnostic performance. A) Receiver operating characteristic (ROC) analyses of the SARS-CoV-2 ELISA using the RBD domain as antigen and 220 negative animal sera (ferret, raccoon dog, cat, cattle, chicken, rabbit and hamster) and 59 sera of infected or vaccinated animals. B) Reproducibility and repeatability of the ELISA. A negative and a very weak positive cattle serum, as well as a positive sample collected from a raccoon dog were tested in five replicates each in five independent approaches. The box plots represent the results of all 25 respective replicates. Each outlier is marked by a circle. C) Performance of the SARS-CoV-2 ELISA for animal sera from clinical trials. Samples taken on day 8 or 12 after infection are shown by open squares. Sera that were collected from day 15 onwards after an experimental infection (ferret, raccoon dog, cattle, hamster) or immunization (chicken, rabbit), or after the first RT-PCR positive throat swab sample (cat) are indicated by filled squares. Negative control samples are shown by circles. The total number of sera per group is given in brackets.

When assessing different sample and conjugate concentrations on the RBD-coated plates, an optimized signal-to-noise ratio was achieved by using serum and multi-species conjugate dilutions of 1/100 and 1/80, respectively (figure 2). Hence, all following analyses were carried out using this test setup.

In order to evaluate both the sensitivity and specificity of the final test protocol and to establish a threshold for positivity, SARS-CoV-2 antibody-negative and -positive sera of multiple species were tested. ROC curve analyses indicated that the ELISA had a very high diagnostic accuracy with an area under the curve (AUC) of 0.989 (95% confidence interval (CI): 0.969 to 1.00). Based on the ROC curve (figure 3A), a cut-off of ≤0.2 for negativity and ≥0.3 for positivity was set, with the intermediate zone between 0.2 and 0.3 being inconclusive. Using these cut-off values, an overall diagnostic specificity of 100.0% (95% CI: 98.34% to 100.0%) and sensitivity of 98.31% (95% CI: 90.91% to 99.96%) was achieved, where the optimal specificity is particularly remarkable, as in some of the animal sera high titers of antibodies against non-SARS beta-coronaviruses were present like bovine coronaviruses (Ulrich, Wernike, Hoffmann, Mettenleiter, & Beer, 2020). Consequently, the previously described high suitability and accuracy of RBD-based antibody detection systems for human coronaviruses (Chia et al., 2020; Meyer et al., 2014) could also be confirmed for different animal species.

Regarding the diagnostic sensitivity, every antibody positive serum included in this study could be identified, with only one exception (figure 3C). This was a single putative seropositive sample that was collected from a ferret 12 days after experimental infection, which tested negative by VNT and exhibited a low titer of 1/64 in the iIFA (ferret number 5 described in (Schlottau et al., 2020)). When considering only sera collected later than 12 days after infection, every individual was correctly detected as being antibody-positive, independent of the animal species. Thus, the newly established ELISA protocol enables the antibody detection in a broad range of animal species that are susceptible to SARS-CoV-2, such as ferrets, cats, raccoon dogs, hamsters or cattle. Furthermore, the applicability could be shown for further species like rabbits or chicken (figure 3C), which are frequently used for reagent production for research, diagnostics and therapeutic purposes (Farzaneh, Hassani, Mozdziak, & Baharvand, 2017; Popkov et al., 2003; Rossi et al., 2005). Hence, the versatility was demonstrated for a test system that does not require high-containment laboratories as e.g. species-independent neutralization assays. However, when further species not included in the current validation are to be tested, the test has to be validated including the suitability of the conjugate for the particular species in question. Alternatively, a species-specific conjugate may be used, as shown here for chicken (figure 3).

In the final step of the validation, both the repeatability and reproducibility were assessed using five replicates each of three sera in five independent ELISA assays. The negative cattle serum and the positive raccoon dog sample tested correctly in each case with very low variations (figure 3B). Mean OD values and standard deviations of 0.03±0.02 and 0.72±0.07 were calculated for the antibody-negative and -positive sample, respectively. Every replicate of the very weak positive cattle sample tested positive or resulted in the doubtful measuring range of the ELISA (mean 0.28, standard deviation 0.03, min 0.21, max 0.33). Thus, the animal would have been detected as SARS-CoV-2-antibody positive in every approach (figure 3B), further demonstrating the reliability of the test system.

In conclusion, we established a versatile ELISA protocol for the highly sensitive and specific detection of SARS-CoV-2 antibodies in animal sera, which could be used for outbreak investigations, monitoring studies or to identify yet unknown reservoir or intermediate hosts.

## Acknowledgments

We thank Bianka Hillmann and Mareen Lange for excellent technical assistance. This research was supported by intramural funding of the German Federal Ministry of Food and Agriculture provided to the Friedrich-Loeffler-Institut.

## Data availability statement

The data that support the findings of this study are available from the corresponding author upon reasonable request.

## Ethical Statement

The antibody negative sera represented routine diagnostic submissions taken by the responsible veterinarians in the context of health monitoring (no permissions were necessary to collect the specimens), or originated from unrelated animal trials, which were reviewed by the responsible state ethics commission and were approved by the competent authority (State Office of Agriculture, Food Safety, and Fishery in Mecklenburg-Western Pomerania). The SARS-CoV-2 antibody positive specimens were collected in the context of infection trials, monitoring of the natural course of infection, or immunizations conducted for diagnostic reagent production (permission numbers MV/TSD/7221.3-2-010/18-12 and 7221.3-2-042/17 (State Office of Agriculture, Food Safety, and Fisheries in Mecklenburg-Western Pomerania), 33.8-42502-05-20A522 (Niedersächsisches Landesamt für Verbraucherschutz und Lebensmittelsicherheit) and 0086/20 (Landesamt für Gesundheit und Soziales in Berlin)).

